# Inactivation of prenyltransferase inhibitors by S-methylation catalyzed by the methyltransferases TMT1A (METTL7A) and TMT1B (METTL7B)

**DOI:** 10.64898/2026.11.12.766183

**Authors:** Diya V. Jariwala, Lucia A. Lavallee, Michelle A. Kwarteng, William J.E. Frye, Zachary E. Paikin, Connor P. Jewell, Lisa M. Jenkins, Crystal C. Lipsey, Robert W. Robey, Michael M. Gottesman, Pedro J. Batista

**Author notes:** Equal contribution (Diya V. Jariwala, Lucia A. Lavallee). Correspondence should be addressed to P.J.B., 37 Convent Drive, Building 37, Room 2128, Bethesda, MD, 20892.

## Abstract

The thiol methyltransferases TMT1A and TMT1B (formerly METTL7A and METTL7B) metabolize thiol-containing compounds, but their substrate range and effects on drug activity remain incompletely defined. Here, we investigated whether these enzymes methylate protein prenyltransferase inhibitors and alter cellular sensitivity to these compounds. Mass spectrometry identified several thiol-containing farnesyltransferase and geranylgeranyltransferase inhibitors as substrates of both enzymes, whereas other thiol-containing inhibitors showed no detectable methylation. In HT1080 fibrosarcoma cells, TMT1A or TMT1B overexpression conferred resistance to several geranylgeranyltransferase inhibitors and attenuated drug-induced cell-cycle arrest. Pharmacological inhibition of TMT1A with 2,3-Dichloro-α-methylbenzylamine (DCMB, LY-78335) reversed resistance and restored cell-cycle arrest in TMT1A-expressing cells. These findings expand the known substrate range of TMT1A and TMT1B and support TMT-dependent methylation as a mechanism that reduces the cellular activity of susceptible prenyltransferase inhibitors.

## Introduction

The human genome encodes approximately 200 methyltransferases (MTases), most of which use S-adenosylmethionine (SAM) as the methyl donor [1, 2]. These enzymes transfer methyl groups to oxygen, nitrogen, carbon, or sulfur atoms in substrates such as proteins, nucleic acids, and small molecules [2]. The MTases that methylate small molecules include catechol *O*-methyltransferase (COMT) and thiopurine *S*-methyltransferase (TPMT). COMT methylates catechol-containing drugs such as L-DOPA, carbidopa, and apomorphine, used in the treatment of Parkinson’s disease [3], whereas TPMT methylates 6-thioguanine and 6-mercaptopurine, used in the treatment of leukemia [4]. The thiol methyltransferases TMT1A and TMT1B (formerly METTL7A and METTL7B, respectively) are recent additions to the family of small-molecule MTases. Both enzymes methylate several sulfur-containing compounds, including hydrogen sulfide, captopril, thiospironolactone, and dithiothreitol [5, 6], but do not methylate endogenous thiols such as glutathione and cysteine. Known substrates of TMT1A and TMT1B include certain histone deacetylase (HDAC) inhibitors and angiotensin-converting enzyme (ACE) inhibitors that contain thiol groups capable of coordinating with the catalytic zinc ion of their respective target enzymes [7, 8]. Methylation of this zinc-binding thiol underlies the resistance to certain thiol-containing HDAC inhibitors conferred by TMT1A or TMT1B overexpression [9]. However, the range of thiol-containing drugs susceptible to TMT1A- and TMT1B-mediated methylation and the consequences for drug resistance remain incompletely defined. These observations prompted us to investigate thiol-containing inhibitors of other zinc-dependent enzymes.

The protein prenyltransferases (PTases), farnesyltransferase (FTase) and geranylgeranyltransferase (GGTase), are also zinc-dependent enzymes that catalyze the covalent attachment of isoprenoid groups to the cysteine residues near the C-terminus of proteins, promoting proper subcellular localization [10]. Among the best-characterized substrates of PTases are members of the RAS family of small GTPases. Both wild-type and oncogenic forms of HRAS, KRAS, and NRAS require prenylation for efficient membrane association and activation of downstream signaling pathways [11]. Although HRAS is primarily farnesylated, KRAS and NRAS can undergo alternative geranylgeranylation by Geranylgeranyltransferase Type-1 (GGTase-1) upon FTase inhibition, providing a rationale for dual inhibition of these enzymes [12]. Early inhibitors of FTase and GGTase-1, including L-744,832 and GGTI-298, were peptidomimetics of the prenylation CAAX consensus motif present in all RAS proteins and contained a thiol group that mimicked the cysteine thiol in this motif [13-15]. Like thiol-containing HDAC and ACE inhibitors, these PTase inhibitors rely on a thiol group for zinc binding and enzyme inhibition [16]. We therefore determined whether these thiol-containing PTase inhibitors are substrates of TMT1A or TMT1B. Here, we show that TMT1A and TMT1B methylate a subset of thiol-containing farnesyltransferase inhibitors (FTIs) and geranylgeranyltransferase inhibitors (GGTIs). In HT1080 cells, whose growth is inhibited by GGTIs [17], overexpression of either enzyme confers resistance to GGTIs and prevents GGTI-induced cell-cycle arrest. Our findings identify thiol methylation as a previously unrecognized mechanism of resistance to protein prenylation inhibitors and expand the known repertoire of TMT1A and TMT1B substrates.

## Materials and methods

### Chemicals

Romidepsin (molecular weight, 540.7 g/mol) was purchased from Selleck Chemicals (Houston, TX). GGTI-298 (molecular weight, 479.63 g/mol), FTI-2153 (molecular weight, 466.6 g/mol), GGTI-2418 (molecular weight, 441.52 g/mol), GGTI-2154 (molecular weight, 420.5 g/mol), FTI-I (molecular weight, 470.7 g/mol), and FTI-II (molecular weight, 371.5 g/mol) were purchased from GLPBIO (Montclair, CA). KD5170 (molecular weight, 451.6 g/mol), NCH-51 (molecular weight, 390.6 g/mol), tipifarnib (molecular weight, 489.4 g/mol), and GGTI-2133 (molecular weight, 456.5 g/mol) were purchased from Cayman Chemical (Ann Arbor, MI). GGTI-286 (molecular weight, 429.58 g/mol) was purchased from AOBIOUS (Gloucester, MA). L-744,832 (molecular weight, 632.70 g/mol) and LY-78335 (2,3-Dichloro-α-methylbenzylamine, DCMB; molecular weight, 226.53 g/mol) were purchased from Santa Cruz Biotechnology (Santa Cruz, CA).

### Cell lines

HEK-293 and HT1080 cells (both from ATCC, Manassas, VA) were cultured in Eagle’s minimum essential medium (EMEM) supplemented with 10% fetal bovine serum (FBS), 2 mM L-glutamine, and 100 U/mL penicillin– streptomycin. Cells were transfected with an empty pcDNA3.1 vector or pcDNA3.1 encoding human TMT1A or TMT1B fused to a C-terminal FLAG tag (GenScript, Piscataway, NJ) using Lipofectamine 2000 (Thermo Fisher Scientific, Waltham, MA) according to the manufacturer’s instructions. Stable single-cell clones were isolated by G418 selection (1 mg/mL; Corning Mediatech, Manassas, VA) and maintained under continuous selection. HEK-293 cells expressing empty vector, TMT1A, or TMT1B have been previously described [9].

### Mass spectrometry

HEK-293 cells transfected with empty vector, or transfected to express TMT1A or TMT1B, were plated in 6-well plates and allowed to attach overnight. PTase inhibitors (10 µM) were then added to the cells and incubated for 6 hours, after which the culture medium was collected. Culture medium was centrifuged at 15000 × g for 15 minutes to pellet insoluble material and 50 μL of supernatant was removed for mass spectrometry analysis. For each sample, 10 μL was injected into an Exion liquid chromatography system (SCIEX, Framingham, MA) coupled inline to an X500B QTOF mass spectrometer (SCIEX, Framingham, MA). Each sample was separated on a 2.1 x 50 mm Aeris 3.6 μm WIDEPORE XB-C8 200 Å column (Phenomenex, Torrance, CA) at 0.3 mL/min using a 4 min linear gradient from 2% to 40% solvent B in solvent A (A: 0.1% formic acid; B: acetonitrile containing 0.1% formic acid). Data were acquired in positive ion mode over a mass range of 100-800 m/z with data-dependent MS/MS acquisition performed over the mass range of 50-750 m/z. Relative quantitation was performed from extracted ion chromatogram peak areas using SCIEX OS software (SCIEX, Framingham, MA, RRID:SCR_023651). Percentage methylation was calculated as 100 × methylated peak area/(parent + methylated peak areas).

### Cytotoxicity assays

Cells were seeded in white, opaque 96-well plates at 5,000 cells per well and allowed to attach overnight. Outer wells were filled with sterile phosphate-buffered saline (PBS) to minimize edge effects caused by evaporation. Cells were subsequently treated with the indicated compounds in triplicate and incubated for an additional 72 h. Cell viability was then measured using the CellTiter-Glo Luminescent Cell Viability Assay (Promega, Madison, WI) according to the manufacturer’s instructions. Luminescence was measured using an Infinite M200 PRO microplate reader (Tecan, RRID:SCR_019033, San Jose, CA). Dose-response curves were generated in GraphPad Prism (GraphPad Software), and half-maximal inhibitory concentrations (IC_50_) were calculated by nonlinear regression using a four-parameter logistic model.

### Western blot

Whole-cell lysates were prepared by lysing cells in RIPA buffer (50 mM Tris-HCl [pH 7.4], 1% Triton X-100, 10% glycerol, 0.1% SDS, 2 mM EDTA, and 0.5% deoxycholate, 50 mM NaCl) with protease inhibitor cocktail (cat# 5871, Cell Signaling Technology, Danvers, MA) and sonicating using an inverted cup-horn sonicator (three 30 s bursts). Lysates were clarified by centrifugation at 10,000 × g for 10 min, and the supernatants were collected. Protein concentrations were determined using the Bio-Rad Protein Assay (Bio-Rad Laboratories, Hercules, CA). Approximately 25 μg of total protein per sample was resolved by SDS-PAGE and transferred overnight onto nitrocellulose membranes. Membranes were probed with antibodies against FLAG (Sigma-Aldrich, St. Louis, MO; catalog no. F1804-200UG), TMT1A (Proteintech Group, Rosemont IL; catalog no. 17092-1-AP), TMT1B (Proteintech Group; catalog no. 17001-1-AP), and β-actin (Cell Signaling Technology; catalog no. 3700), followed by IRDye-conjugated secondary antibodies (LI-COR Biosciences, Lincoln, NE). Immunoreactive bands were detected using an Odyssey CLx Imaging System (LI-COR Biosciences).

### Cell cycle analysis

Cell cycle experiments were performed as previously described [9]. Briefly, cells were plated and allowed to attach overnight after which the desired concentration of PTase inhibitor was added for a 24 h incubation. Cells were then harvested by trypsinization, washed, and resuspended in equal volumes of staining solution (50 µg/ml propidium iodide and 0.1% Triton-X in DI water) and RNAse solution (200 U/ml RNase in DI water). After a 20 min incubation, propidium iodide fluorescence was detected using a FACS CANTO II flow cytometer equipped with a 488 nm argon laser. At least 10,000 events were collected per sample. The percentage of cells in each phase of the cell cycle was determined using ModFitLT 5.0 software (Verity Software House, Topsham, ME).

## RESULTS

### TMT1A and TMT1B methylate thiol-containing FTIs and GGTIs

To determine whether the reactive thiols in farnesyltransferase inhibitors (FTIs) and geranylgeranyltransferase inhibitors (GGTIs) are susceptible to enzymatic thiol methylation by TMT1A or TMT1B, we treated HEK-293 cells expressing either enzyme and matched empty-vector control cells with a panel of these inhibitors. We chose HEK-293 cells because their lack of sensitivity to these drugs allowed us to assess drug methylation without the confounding effects of reduced cell fitness. We and others have previously shown that methylated forms of compounds such as captopril, thiospironolactone, and romidepsin can be detected in the culture medium after incubation with cells expressing TMT1A or TMT1B [6, 9, 18]. HEK-293 cells expressing TMT1A or TMT1B were incubated with KD5170 and NCH-51 at 10 µM for 6 hours as positive controls. As previously observed, methylated KD5170 and NCH-51 levels were higher in the culture medium of enzyme-expressing cells (Supplemental Figure 1A). In agreement with the observation that cells expressing TMT1B were less resistant to KD5170 in cytotoxicity assays [9], we detected slightly less methylated drug in cells expressing TMT1B. Under the same conditions, we tested a panel of PTase inhibitors. Significantly higher levels of methylated GGTI-298, GGTI-286, and L-744,832 were detected in the culture medium of cells expressing either enzyme than in that of empty-vector controls (Figure 1A, Supplemental Table 1). All three compounds contain a thiol group (Supplemental Figure 2). No methylation was detected for the thiol-containing FTIs FTI-I and FTI-II or for the negative-control FTase inhibitor tipifarnib. Together, these findings demonstrate that the presence of a thiol group alone is not sufficient for TMT1A- or TMT1B-dependent methylation and identify a subset of thiol-containing PTase inhibitors as substrates of both enzymes.

**Figure 1:**
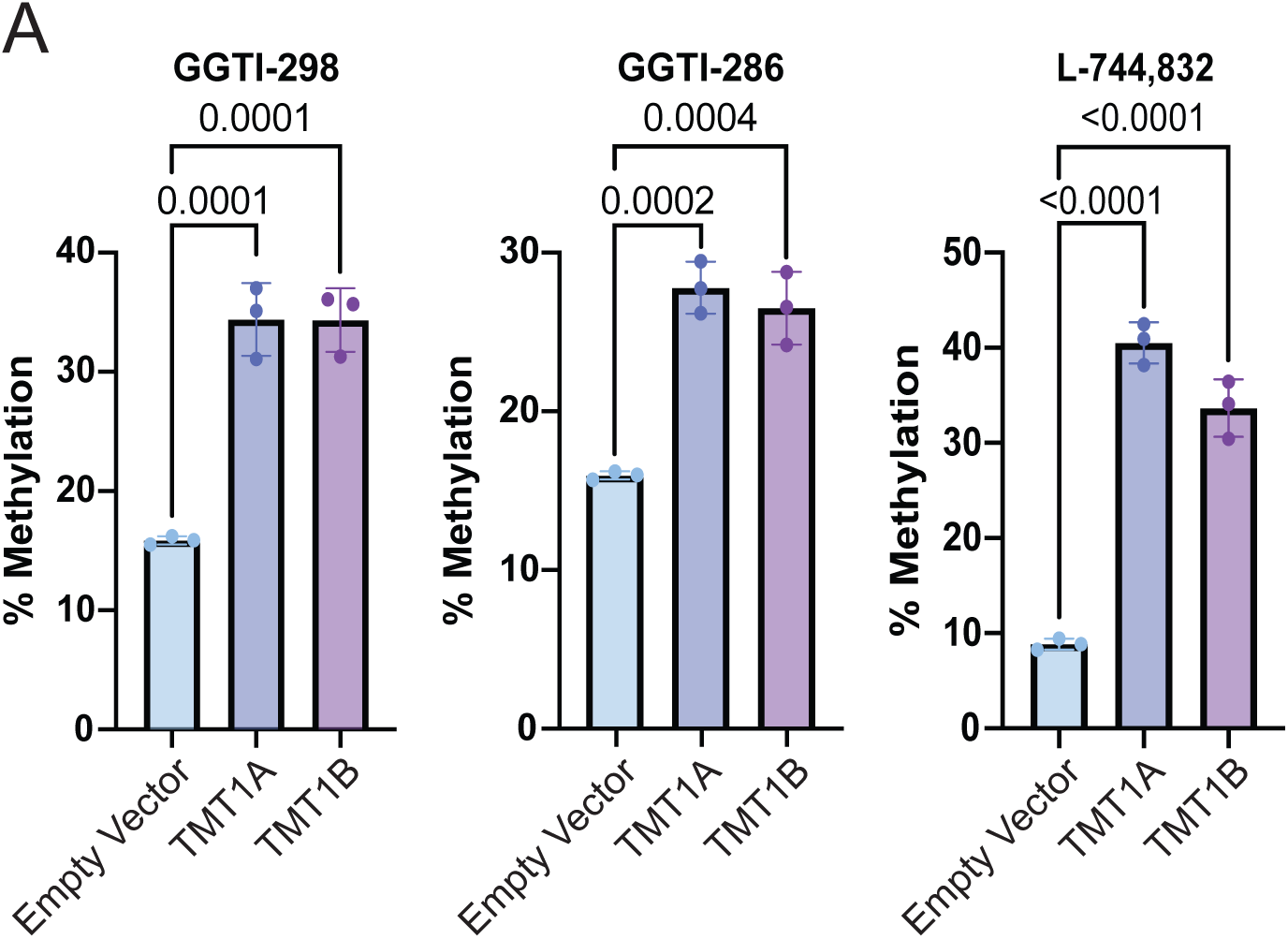
**(A)** Measurement of methylated drug by liquid chromatography– tandem mass spectrometry (LC–MS/MS) in cell media following 6 hours of incubation with HEK-293 cells expressing TMT1A or TMT1B. Error bars represent mean ± standard deviation (n=3 biological replicates). Dots represent individual measurements. One-way ANOVA p-values are displayed.

### Establishment and validation of the HT1080 cell model

Having identified PTase inhibitors methylated by TMT1A and TMT1B, we next examined whether TMT-dependent methylation is associated with reduced inhibitor activity and resistance to these compounds. To address this question, we generated HT1080 fibrosarcoma cells stably expressing either TMT1A or TMT1B. HT1080 cells lack detectable endogenous TMT1A and TMT1B expression (Figure 2A) and have previously been shown to undergo cell-cycle arrest in response to PTase inhibitors such as GGTI-298 [17]. We examined TMT1A or TMT1B expression using protein-specific antibodies and an anti-FLAG antibody that detects both exogenously expressed proteins. We found that HT1080 cells express TMT1A and TMT1B at levels comparable to those of the transfected HEK-293 cells (Figure 2A) used above for the mass spectrometry analysis. To confirm that overexpressed TMT1A and TMT1B were functional in HT1080 cells, we treated the cells with the thiol-containing HDAC inhibitors KD5170, NCH-51, and romidepsin, which we previously identified as substrates of TMT1A and/or TMT1B [9], as well as a non-thiol HDAC inhibitor, vorinostat. Overexpression of either enzyme conferred similar levels of resistance to NCH-51, whereas resistance to KD5170 was lower in cells expressing TMT1B than in those expressing TMT1A. Appreciable resistance to romidepsin was observed only in cells expressing TMT1A, and as expected, expression of TMT1A or TMT1B had no effect on sensitivity to vorinostat, consistent with our previous results [17] (Figure 2B, Supplemental Table 2).

**Figure 2:**
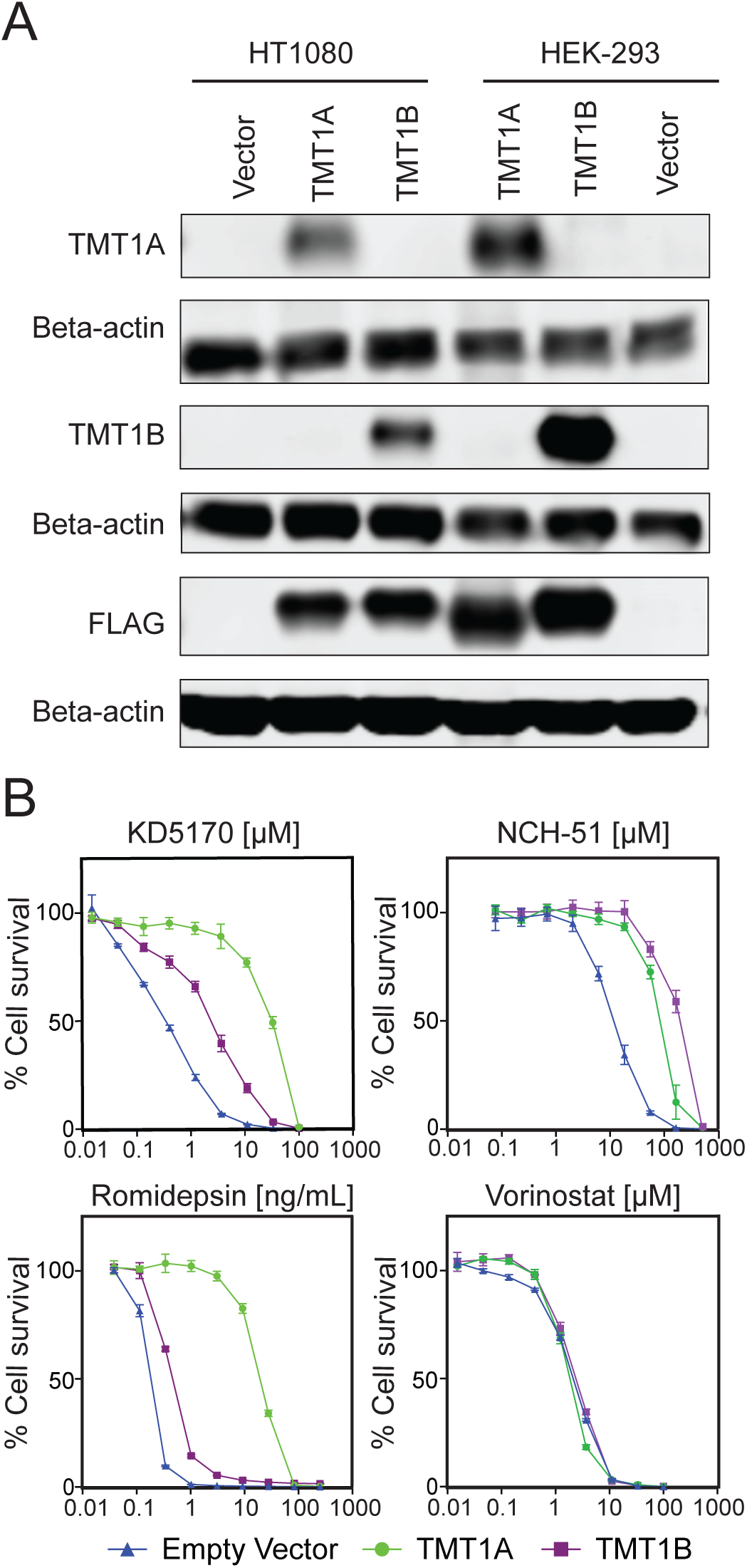
**(A)** Western blot analysis of TMT1A and TMT1B expression in HT1080 and HEK-293 cells. Blots were probed with antibodies against TMT1A, TMT1B and FLAG. Beta-actin served as a loading control. **(B)** Three-day cytotoxicity assays with KD5170, NCH-51, romidepsin and vorinostat in HT1080 cells expressing TMT1A (circles), TMT1B (squares), or carrying an empty vector (triangles). Error bars represent mean ± standard deviation (n=3 biological replicates).

### TMT1A and TMT1B overexpression confers resistance to a subset of PTase inhibitors

We next examined whether expression of TMT1A or TMT1B in HT1080 cells could alter sensitivity to PTase inhibitors that can be methylated by TMT1A or TMT1B. Among the GGTase inhibitors tested, overexpression of either TMT1A or TMT1B conferred resistance to the thiol-containing inhibitor GGTI-298, consistent with our mass spectrometry data showing increased methylation in cells expressing either enzyme (Figure 3A, Supplemental Table 2). Overexpression of either enzyme also conferred modest resistance to the thiol-containing inhibitor GGTI-286 relative to empty-vector controls (Figure 3A, Supplemental Table 2). In contrast, neither enzyme altered sensitivity to GGTI-2418, GGTI-2133, or GGTI-2154, which lack thiol groups (Supplemental Figure 3A, Supplemental Table 2).

**Figure 3:**
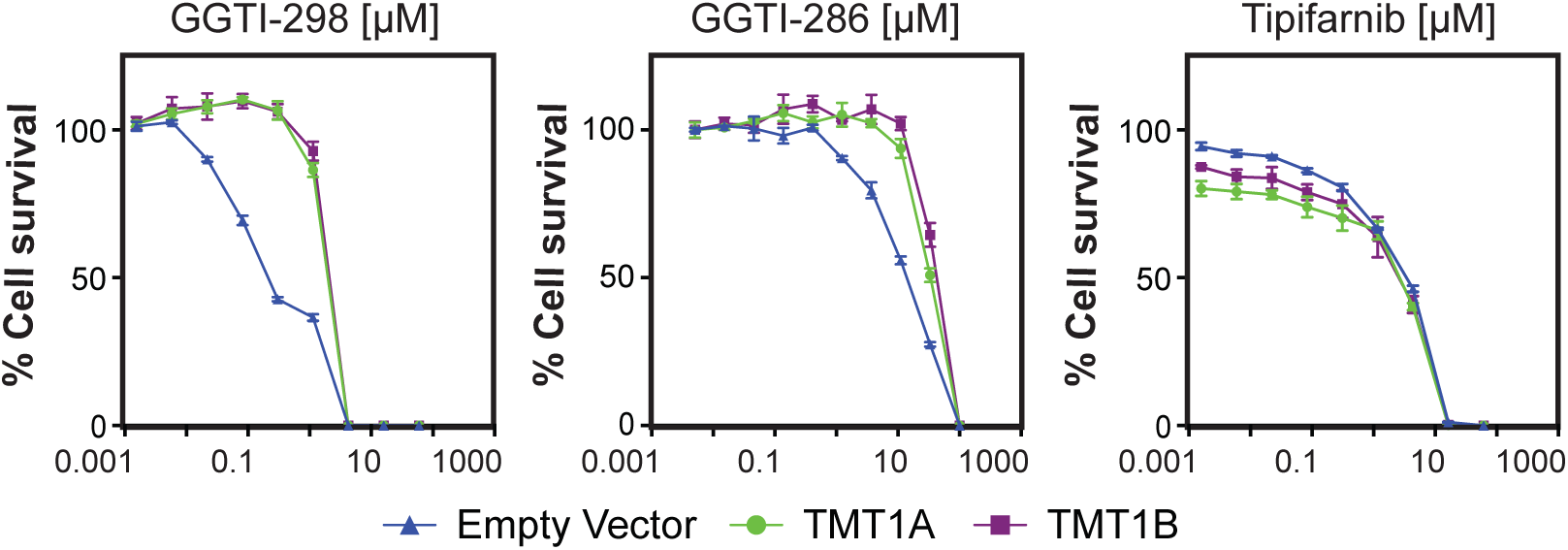
Three-day cytotoxicity assays with **(A)** GGTI-298 and GGTI-286 or **(B)** tipifarnib in HT1080 cells expressing TMT1A (circles), TMT1B (squares), or carrying an empty vector (triangles). Error bars represent mean ± standard deviation (n=3 biological replicates).

Among the FTIs examined, tipifarnib inhibited the growth of HT1080 empty-vector control cells. Expression of TMT1A or TMT1B did not alter sensitivity to tipifarnib, consistent with its lack of a thiol group (Figure 3B, Supplemental Table 2). While we found that L-744,832 could be methylated by TMT1A and TMT1B, L-744,832 showed relatively low potency in HT1080 cells, limiting our ability to assess whether overexpression of either enzyme conferred resistance. FTI-I, FTI-II, and FTI-2153 also showed relatively low potency in HT1080 cells (Supplemental Figure 3B, Supplemental Table 2). Thus, we identified a subset of PTase inhibitors for which overexpression of TMT1A or TMT1B could methylate and inactivate the drug, thereby conferring resistance.

### Cell-cycle arrest induced by GGTase inhibitors is rescued by TMT1A and TMT1B

GGTI-298 has previously been shown to induce G_0_/G_1_ cell-cycle arrest in HT1080 cells [17]. We therefore asked whether the resistance conferred by TMT1A or TMT1B to GGTI-298 or GGTI-286 was accompanied by protection from GGTI-induced cell-cycle arrest. Treatment of HT1080 EV cells with 10 µM GGTI-298 or 25 µM GGTI-286 for 24 h resulted in an accumulation of cells in G_0_/G_1_ (Figure 4A). In contrast, expression of either TMT1A or TMT1B significantly reduced G_0_/G_1_ accumulation induced by GGTI treatment (Figure 4A). As noted above, HT1080 cells were sensitive to treatment with the non-thiol FTI tipifarnib. Similarly, treatment of HT1080 EV cells with 10 µM tipifarnib for 24 h resulted in a G_0_/G_1_ cell-cycle arrest, yet overexpression of TMT1A or TMT1B did not protect cells, consistent with the fact that tipifarnib does not have a thiol. Together, these findings demonstrate that TMT1A and TMT1B protect HT1080 cells from G_0_/G_1_ arrest induced by thiol-containing GGTIs and further support a role for TMT enzymatic activity in resistance to these compounds.

**Figure 4:**
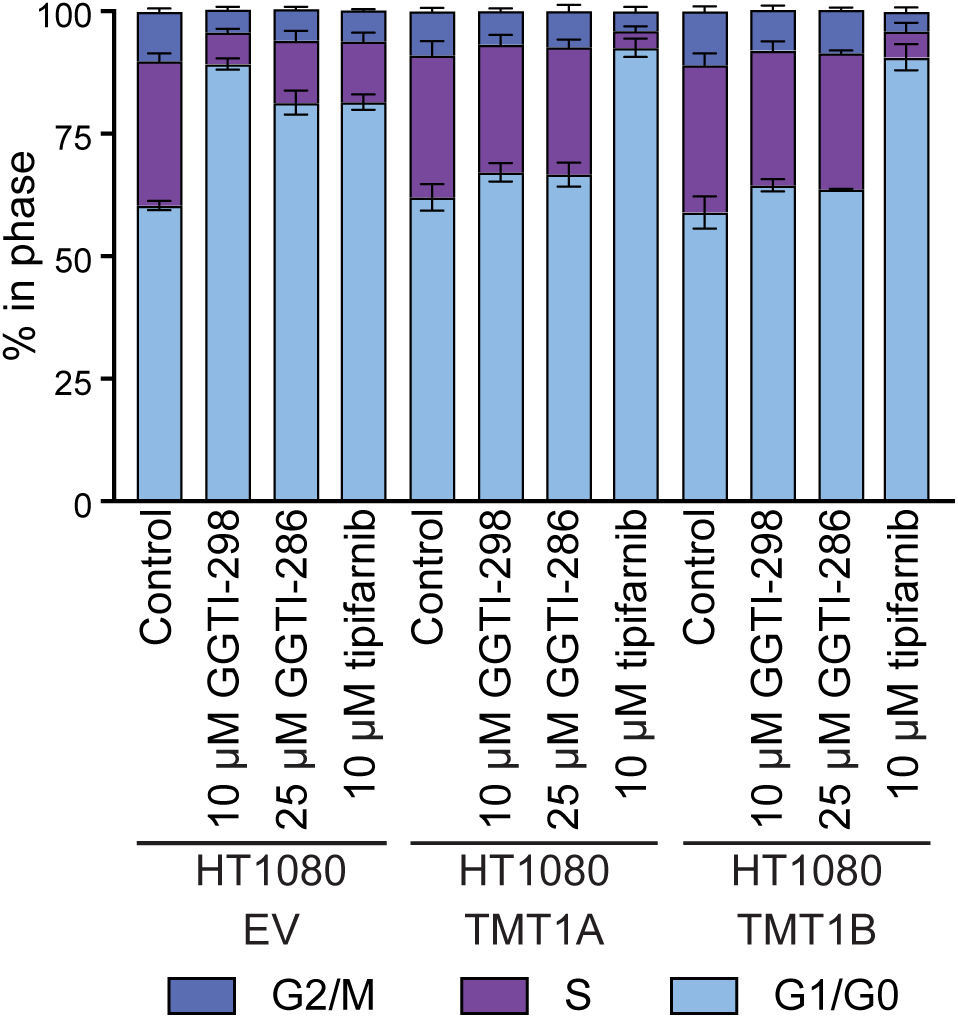
**(A)** Percentage of cells in each phase of the cell cycle for HT1080 EV, HT1080 TMT1A and HT1080 TMT1B cells treated with GGTI-298, GGTI-286, or tipifarnib. Data were compiled from three independent experiments. Error bars indicate standard deviation.

### Inhibition of TMT1A by DCMB restores sensitivity to GGTase inhibitors

DCMB is a phenylethanolamine N-methyltransferase inhibitor that preferentially inhibits TMT1A over TMT1B [6]. To determine whether TMT1A enzymatic activity is required for resistance to the GGTIs examined above, we performed cytotoxicity assays and cell-cycle analyses in the presence or absence of 100 µM DCMB. Pharmacological inhibition of TMT1A with DCMB partially reversed resistance to GGTI-298 and completely reversed resistance to GGTI-286 in HT1080 cells expressing TMT1A, but had no effect in empty-vector controls or cells expressing TMT1B (Figure 5A, Supplemental Table 3). Consistent with these findings, cotreatment with 100 µM DCMB and either 10 µM GGTI-298 or 25 µM GGTI-286 restored G₀/G₁ cell-cycle arrest in cells expressing TMT1A, but had no effect in empty-vector controls or cells expressing TMT1B (Figure 5B). Together with the methylation data, these findings support a model in which TMT1A expression confers resistance to a subset of thiol-containing GGTase inhibitors through inhibitor methylation and inactivation.

**Figure 5:**
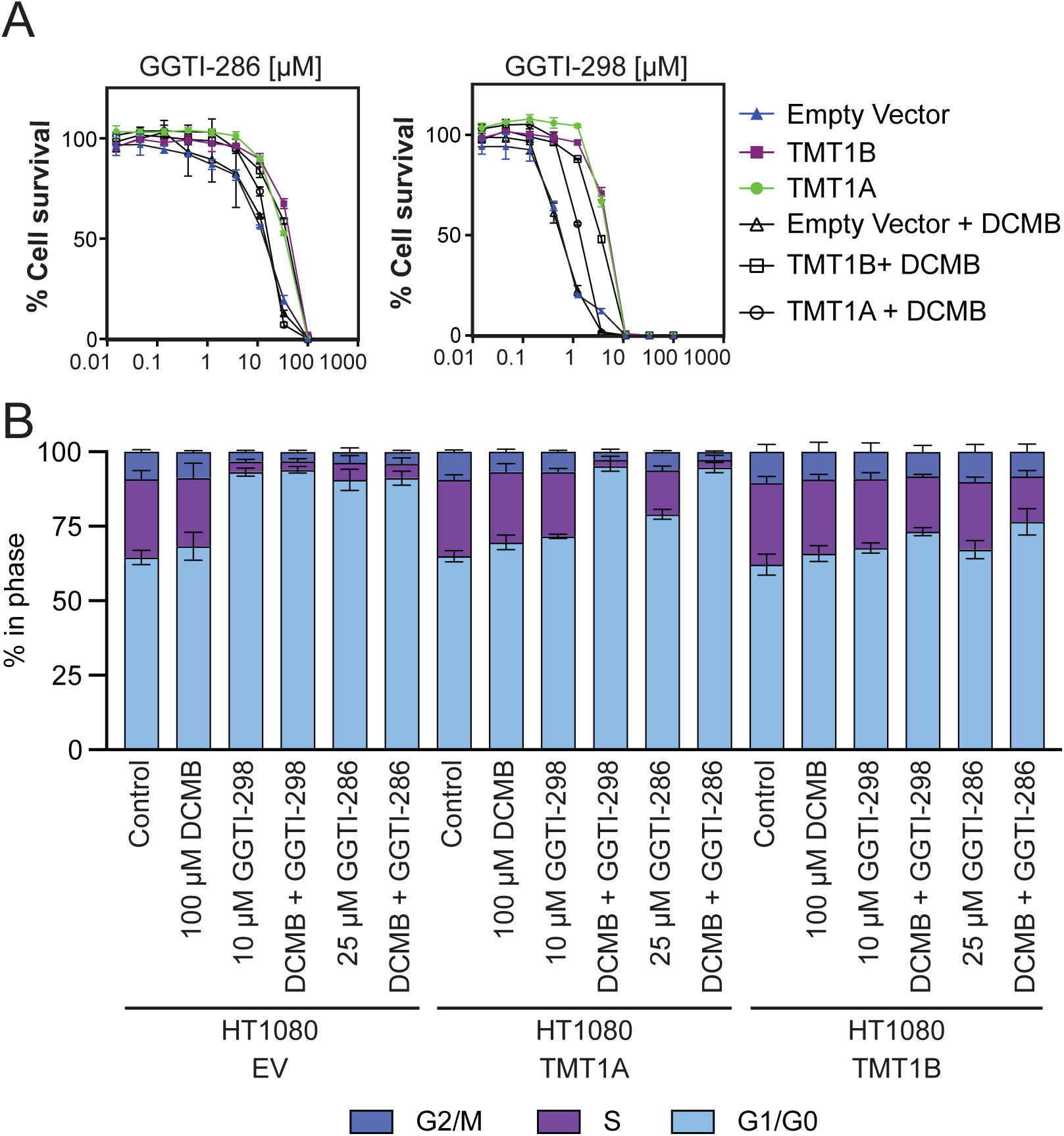
**(A)** Three-day cytotoxicity assays with GGTI-286 and GGTI-298 in the presence or absence of DCMB in HT1080 cells expressing TMT1A (circles), TMT1B (squares), or empty vector (triangles). Open symbols indicate the presence of DCMB; filled symbols indicate its absence. Error bars represent mean ± standard deviation (n=3 biological replicates). **(B)** Percentage of cells in each cell cycle phase for HT1080 EV, HT1080 TMT1A and HT1080 TMT1B cells treated with GGTI-298 or GGTI-286 in the presence or absence of DCMB. Data were compiled from three independent experiments. Error bars indicate standard deviation.

## Discussion

S-methylation plays a significant role in the metabolism of drugs containing thiol-moieties [19]. Although the existence of TMT was postulated decades ago, with dithiothreitol, 2-mercaptoethanol, captopril and thiospironolacone among the earliest recognized substrates [20, 21], the enzymes responsible for this activity were only recently identified as METTL7A and METTL7B, subsequently renamed as TMT1A and TMT1B, respectively [5, 6]. We previously identified TMT1A overexpression as a driver of resistance to the HDACi romidepsin. Mechanistically, TMT1A methylated the drug’s zinc-binding thiol group, thereby inactivating it [9]. Other thiol-containing HDAC inhibitors such as KD5170, NCH-51 and the natural product largazole were also found to be TMT1A and TMT1B substrates [9], leading us to search for other molecules that could be substrates of the enzymes.

Here, we identify thiol-containing peptidomimetic PTase inhibitors as substrates of TMT1A and TMT1B. To determine if PTase inhibitors are substrates of these enzymes in cells, we overexpressed TMT1A or TMT1B in HEK-293 cells, which are not sensitive to PTase inhibitors and allow us to measure TMT1A or TMT1B-dependent drug methylation without the confounding effect related to cell viability. As expected, no TMT-dependent methylation was detected for PTase inhibitors lacking a thiol group. We identified GGTI-298, GGTI-286, and L-744,832 as substrates of both enzymes. However, no TMT-dependent methylation was detected for the thiol-containing PTase inhibitors FTI-I and FTI-II (Supplemental Table 1, Supplemental Figure 2). This result is consistent with the substrate selectivity of TMT1A and TMT1B, which methylate some, but not all, thiol compounds. For example, neither enzyme methylates cysteine or glutathione, endogenous compounds with thiol groups [6]. Early peptidomimetic PTase inhibitors, like FTI-I and FTI-II, are designed to mimic the CAAX motif and incorporate a thiol group corresponding to that of cysteine [13, 16]. The absence of detectable TMT-dependent methylation of FTI-I and FTI-II may therefore reflect structural features shared with cysteine that limit substrate recognition. Further experiments are needed to test this hypothesis.

To determine whether TMT1A- or TMT1B-dependent methylation altered drug activity, we tested these inhibitors in HT1080 cells, which are sensitive to GGTI treatment [17] and lack detectable endogenous expression of either enzyme. This allowed us to generate isogenic cell models expressing TMT1A or TMT1B alongside empty-vector controls lacking detectable TMT1A and TMT1B expression. Expression of either enzyme conferred resistance to the PTase inhibitors GGTI-298 and GGTI-286 identified as substrates in our methylation assays. The relatively low potency of L-744,832 in these cells, with an IC_50_ of 44 µM in empty-vector control cells, limits our ability to determine if TMT-dependent methylation of this drug affects its activity.

Mechanistically, treatment of HT1080 cells with GGTI-298 has been shown to reduce HT1080 cell growth by inducing cell-cycle arrest via inhibition of prenylation of essential small GTPases [17]. Overexpression of TMT1A or TMT1B in HT1080 cells prevented the cell-cycle arrest induced by GGTI-298 and GGTI-286, supporting a role for these methyltransferases in conferring resistance to both drugs. Inhibition of TMT1A with DCMB restored drug-induced cell-cycle arrest, indicating that TMT1A methyltransferase activity contributes to resistance. Together, these findings suggest that TMT1A- or TMT1B-dependent methylation reduces the activity of selected PTase inhibitors, expanding the known substrate range of these enzymes and supporting their role in the metabolism of compounds containing free thiols.

Pharmacologic resistance to anticancer drugs can arise through mechanisms intrinsic to tumor cells, such as altered drug metabolism, sequestration, or efflux, or through cell-extrinsic factors that affect drug delivery and cellular responses within the tumor microenvironment [22]. Our findings show that TMT-dependent methylation may influence the pharmacokinetics of susceptible prenyltransferase inhibitors. TMT1A and TMT1B are expressed in human liver, where they account for much of the microsomal thiol S-methyltransferase activity [6]. Hepatic methylation could therefore reduce exposure to the active, unmethylated inhibitor and limit its delivery to tumor tissue, even in tumors with low TMT expression. This would represent an additional potential barrier to drug activity beyond the cellular resistance observed in our models. Studies in human liver microsomes or hepatocytes, followed by pharmacokinetic analyses of parent compounds and methylated metabolites in plasma and tumor tissue, could establish whether methylation at endogenous enzyme levels limits exposure to active inhibitor in vivo.

The use of thiol groups in drug design presents several challenges, including metabolic instability and potential off-target effects [23, 24]. Our findings identify TMT-dependent methylation as an additional liability for susceptible thiol-containing drugs, highlighting the importance of understanding the molecular determinants of TMT substrate selectivity.

## Acknowledgments

We thank members of Batista and Gottesman laboratories and Dr. Nicole Y. Morgan for critical reading of the manuscript. This research was supported by the Intramural Research Program of the National Institutes of Health (NIH). The contributions of the NIH author(s) are considered Works of the United States Government. The findings and conclusions presented in this paper are those of the author(s) and do not necessarily reflect the views of the NIH or the U.S. Department of Health and Human Services. Mention of trade names, commercial products, or organizations does not imply endorsement by the U.S. Government.

## Author contributions

[R.W.R.], [M.M.G.] and [P.J.B.] conceived and supervised the study. [D.V.J.], [L.A.L.], [M.K.], [C.C.L.], [C.P.J.], [L.M.J.], [W.J.E.F.], [Z.E.P.] performed experiments and analyzed the data. [D.V.J.], [L.A.L.], [R.W.R.] and [P.J.B.] wrote the manuscript, with input from all authors. All authors reviewed and approved the final manuscript.

## Figure Legends

**Supplemental Figure 1:**
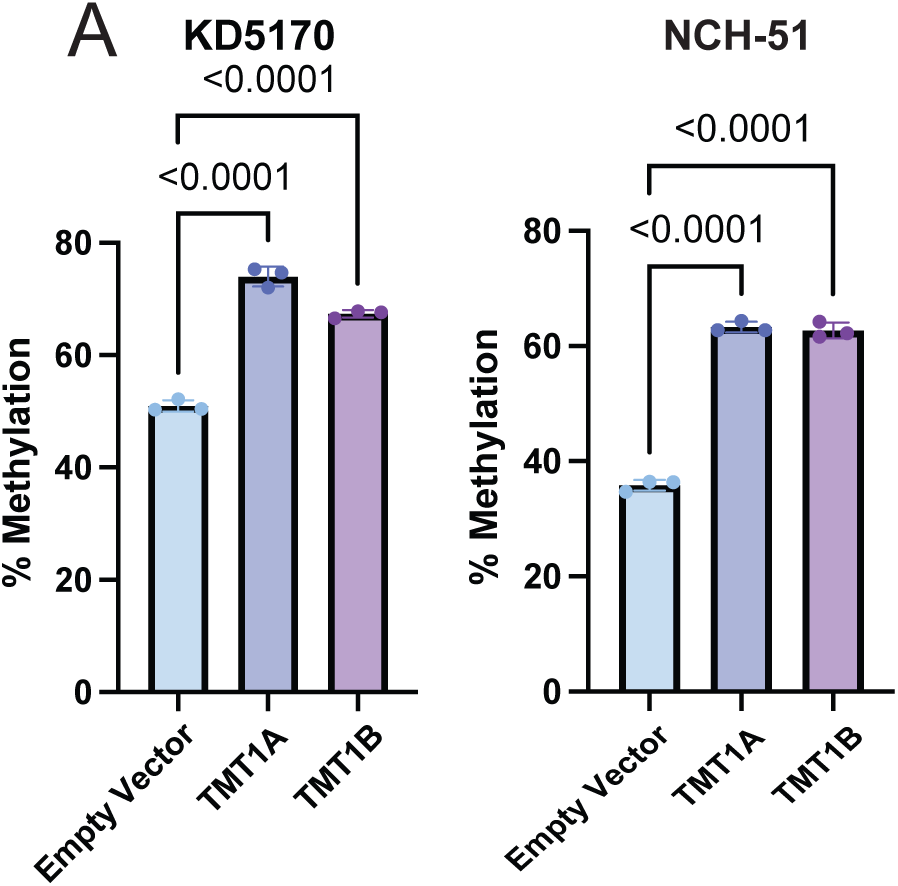
Measurement of methylated HDAC inhibitor by liquid chromatography–tandem mass spectrometry (LC–MS/MS) in cell media following 6 hours of incubation with HEK-293 cells expressing TMT1A or TMT1B. Error bars represent mean ± standard deviation (n=3 biological replicates). Dots represent individual measurements. One-way ANOVA p-values are displayed.

**Supplemental Figure 2:**
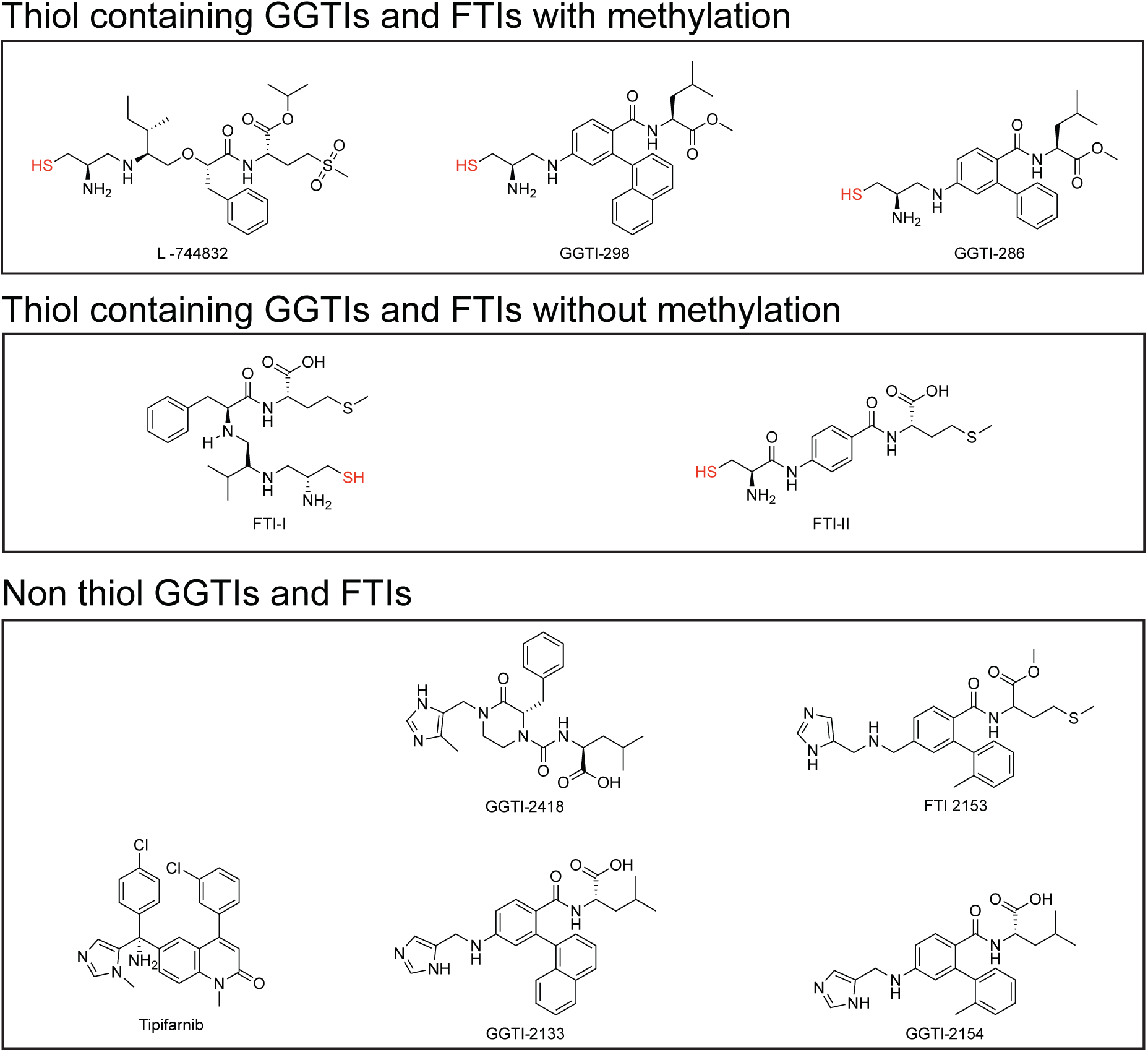
Molecular structures of FTIs and GGTIs used in this study.

**Supplemental Figure 3:**
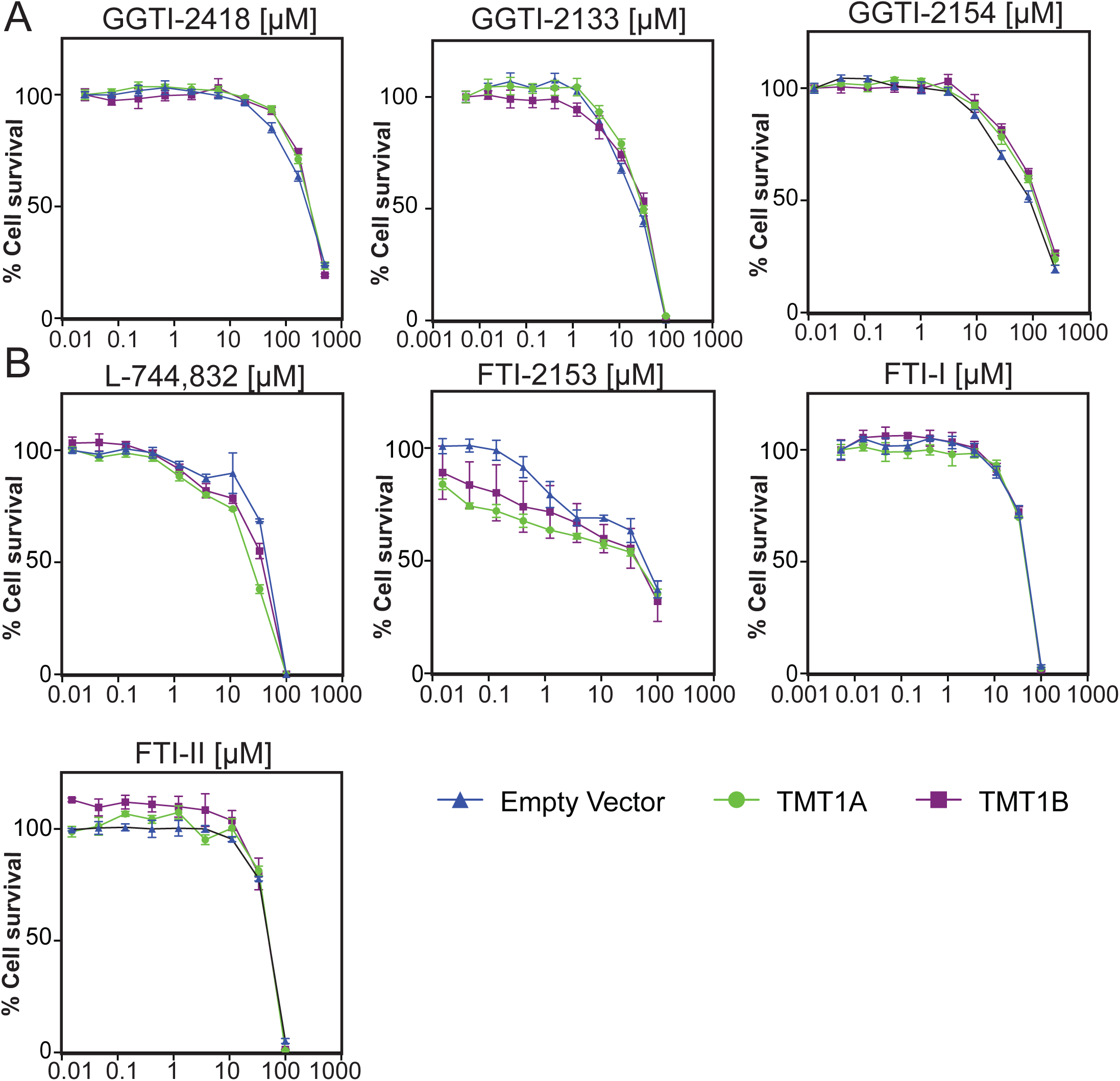
Three-day cytotoxicity assays with (A) GGTI-2418, GGTI-2133 and GGTI-2154 or **(B)** L-744,832, FTI-I, FTI-II or FTI-2153 in HT1080 cells expressing TMT1A (circles), TMT1B (squares), or carrying an empty vector (triangles). Error bars represent mean ± standard deviation (n=3 biological replicates).

**Supplemental Table 1:** FTIs and GGTIs tested for methylation in media of HEK-293 cells expressing TMT1A, TMT1B or carrying an empty vector.

**Supplemental Table 2:** The cross-resistance profile of HT1080 EV, HT1080 TMT1A, and HT1080 TMT1B cells with FTIs and GGTIs.

**Supplemental Table 3:** The cross-resistance profile of HT1080 EV, HT1080 TMT1A, and HT1080 TMT1B cells with thiol-based GGTIs and DCMB.

## Declaration of interest statement

The authors declare no conflicts of interest.

## Notes

### Competing Interest Statement

The authors have declared no competing interest.

## References

[1] A.I. Katsafadou, P.O. Falnes, E. Bruford, B. Braschi, Q.M. Hanson, M.D. Hall, D.C. Thompson, D.W. Nebert, V. Vasiliou, Update of the Methyltransferase Gene Family: Classification, Evolution and Biological Functions, Hum Genomics 20(1) (2026).

[2] E. Abdelraheem, B. Thair, R.F. Varela, E. Jockmann, D. Popadic, H.C. Hailes, J.M. Ward, A.M. Iribarren, E.S. Lewkowicz, J.N. Andexer, P.L. Hagedoorn, U. Hanefeld, Methyltransferases: Functions and Applications, Chembiochem 23(18) (2022) e202200212.

[3] P. Bastos, T. Gomes, L. Ribeiro, Catechol-O-Methyltransferase (COMT): An Update on Its Role in Cancer, Neurological and Cardiovascular Diseases, Rev Physiol Biochem Pharmacol 173 (2017) 1–39.

[4] S. Pathania, R. Bhatia, A. Baldi, R. Singh, R.K. Rawal, Drug metabolizing enzymes and their inhibitors’ role in cancer resistance, Biomed Pharmacother 105 (2018) 53–65.

[5] B.J. Maldonato, D.A. Russell, R.A. Totah, Human METTL7B is an alkyl thiol methyltransferase that metabolizes hydrogen sulfide and captopril, Sci Rep 11(1) (2021) 4857.

[6] D.A. Russell, M.K. Chau, Y. Shi, I.N. Levasseur, B.J. Maldonato, R.A. Totah, METTL7A (TMT1A) and METTL7B (TMT1B) Are Responsible for Alkyl S-Thiol Methyl Transferase Activity in Liver, Drug Metab Dispos 51(8) (2023) 1024–1034.

[7] M. Manal, M.J. Chandrasekar, J. Gomathi Priya, M.J. Nanjan, Inhibitors of histone deacetylase as antitumor agents: A critical review, Bioorg Chem 67 (2016) 18–42.

[8] K.R. Acharya, E.D. Sturrock, J.F. Riordan, M.R. Ehlers, Ace revisited: a new target for structure-based drug design, Nat Rev Drug Discov 2(11) (2003) 891–902.

[9] R.W. Robey, C.M. Fitzsimmons, W.M. Guiblet, W.J.E. Frye, J.M. Gonzalez Dalmasy, L. Wang, D.A. Russell, L.M. Huff, A.J. Perciaccante, F. Ali-Rahmani, C.C. Lipsey, H.M. Wade, A.V. Mitchell, S.S. Maligireddy, D. Terrero, D. Butcher, E.F. Edmondson, L.M. Jenkins, T. Nikitina, V.B. Zhurkin, A.K. Tiwari, A.D. Piscopio, R.A. Totah, S.E. Bates, H.E. Arda, M.M. Gottesman, P.J. Batista, The Methyltransferases METTL7A and METTL7B Confer Resistance to Thiol-Based Histone Deacetylase Inhibitors, Mol Cancer Ther 23(4) (2024) 464–477.

[10] P.J. Casey, M.C. Seabra, Protein prenyltransferases, J Biol Chem 271(10) (1996) 5289–92.

[11] K. Kato, A.D. Cox, M.M. Hisaka, S.M. Graham, J.E. Buss, C.J. Der, Isoprenoid addition to Ras protein is the critical modification for its membrane association and transforming activity, Proc Natl Acad Sci U S A 89(14) (1992) 6403–7.

[12] F.L. Zhang, P. Kirschmeier, D. Carr, L. James, R.W. Bond, L. Wang, R. Patton, W.T. Windsor, R. Syto, R. Zhang, W.R. Bishop, Characterization of Ha-ras, N-ras, Ki-Ras4A, and Ki-Ras4B as in vitro substrates for farnesyl protein transferase and geranylgeranyl protein transferase type I, J Biol Chem 272(15) (1997) 10232-9.

[13] J. Sun, Y. Qian, A.D. Hamilton, S.M. Sebti, Ras CAAX peptidomimetic FTI 276 selectively blocks tumor growth in nude mice of a human lung carcinoma with K-Ras mutation and p53 deletion, Cancer Res 55(19) (1995) 4243–7.

[14] N.E. Kohl, C.A. Omer, M.W. Conner, N.J. Anthony, J.P. Davide, S.J. deSolms, E.A. Giuliani, R.P. Gomez, S.L. Graham, K. Hamilton, et al., Inhibition of farnesyltransferase induces regression of mammary and salivary carcinomas in ras transgenic mice, Nat Med 1(8) (1995) 792–7.

[15] T.F. McGuire, Y. Qian, A. Vogt, A.D. Hamilton, S.M. Sebti, Platelet-derived growth factor receptor tyrosine phosphorylation requires protein geranylgeranylation but not farnesylation, J Biol Chem 271(44) (1996) 27402–7.

[16] S.B. Long, P.J. Hancock, A.M. Kral, H.W. Hellinga, L.S. Beese, The crystal structure of human protein farnesyltransferase reveals the basis for inhibition by CaaX tetrapeptides and their mimetics, Proc Natl Acad Sci U S A 98(23) (2001) 12948–53.

[17] A. Vogt, J. Sun, Y. Qian, A.D. Hamilton, S.M. Sebti, The geranylgeranyltransferase-I inhibitor GGTI-298 arrests human tumor cells in G0/G1 and induces p21(WAF1/CIP1/SDI1) in a p53-independent manner, J Biol Chem 272(43) (1997) 27224–9.

[18] J.M. Gonzalez Dalmasy, C.M. Fitzsimmons, W.J.E. Frye, A.J. Perciaccante, C.P. Jewell, L.M. Jenkins, P.J. Batista, R.W. Robey, M.M. Gottesman, The thiol methyltransferase activity of TMT1A (METTL7A) is conserved across species, Chem Biol Interact 394 (2024) 110989.

[19] H. Mu, L. Ye, B. Wang, Detailed resume of S-methyltransferases: Categories, structures, biological functions and research advancements in related pathophysiology and pharmacotherapy, Biochem Pharmacol 226 (2024) 116361.

[20] R.A. Keith, I. Jardine, A. Kerremans, R.M. Weinshilboum, Human erythrocyte membrane thiol methyltransferase. S-methylation of captopril, N-acetylcysteine, and 7 alpha-thio-spirolactone, Drug Metab Dispos 12(6) (1984) 717-24.

[21] R.M. Weinshilboum, S. Sladek, S. Klumpp, Human erythrocyte thiol methyltransferase: radiochemical microassay and biochemical properties, Clin Chim Acta 97(1) (1979) 59–71.

[22] M.M. Gottesman, O. Lavi, M.D. Hall, J.P. Gillet, Toward a Better Understanding of the Complexity of Cancer Drug Resistance, Annu Rev Pharmacol Toxicol 56 (2016) 85–102.

[23] A. Fruhauf, F.J. Meyer-Almes, Non-Hydroxamate Zinc-Binding Groups as Warheads for Histone Deacetylases, Molecules 26(17) (2021).

[24] L. Zhang, J. Zhang, Q. Jiang, L. Zhang, W. Song, Zinc binding groups for histone deacetylase inhibitors, J Enzyme Inhib Med Chem 33(1) (2018) 714–721.

