## Supplemental Tables for "Inactivation of prenyltransferase inhibitors by S-methylation catalyzed by the methyltransferases TMT1A (METTL7A) and TMT1B (METTL7B)"

**Supplemental Table 1: FTIs and GGTIs tested for methylation in media of HEK-293 cells expressing TMT1A, TMT1B or carrying an empty vector.**

| Drug | Empty Vector | TMT1A | TMT1B |
|---|---|---|---|
| GGTI-298 | 16±0.35 | 34±3.0 | 34±2.7 |
| GGTI-286 | 15±0.25 | 27±1.6 | 26±2.2 |
| L-744,832 | 8.8±0.58 | 41±2.2 | 34±3.0 |
| NCH-51 | 36±0.95 | 63±0.95 | 63±1.4 |
| KD 5170 | 51±1.0 | 74±1.7 | 67±0.67 |
| Tipifarnib | ND | ND | ND |
| FTI-I | ND | ND | ND |
| FTI-II | ND | ND | ND |

Results are reported as % methylation ± SD

ND = not detected

**Supplemental Table 2: The cross-resistance profile of HT1080 EV, HT1080 TMT1A, and HT1080 TMT1B cells with FTIs and GGTIs.**

| **Drug** | **Thiol group** | **HT1080 EV** | **HT1080 TMT1A** | **HT1080 TMT1B** | **Units** |
|---|---|---|---|---|---|
| KD 5170 | Yes | 0.77 ±0.13 | 118±3.5 | 67.0±2.1 | μM |
| NCH-51 | Yes | 8.4±3.5 | 84±9.4 | 197±18 | μM |
| FTI-2153 | No | 32±20 | 23±13 | 26±4.8 | μM |
| Tipifarnib | No | 7.0±2.1 | 5.5±2.2 | 6.6±1.4 | μM |
| GGTI-2418 | No | 290±33 | 440±90 | 427±108 | μM |
| GGTI-2133 | No | 25±0.89 | 30±3.7 | 31±4.8 | μM |
| GGTI-2154 | No | 82±14 | 122±39 | 146±63 | μM |
| Vorinostat | No | 2.6±0.67 | 2.4±0.80 | 2.4±1.0 | μM |
| Romidepsin | Yes | 0.27±0.081 | 20±1.6 | 0.48±0.037 | μg/mL |
| GGTI-298 | Yes | 1.1±0.58 | 4.5±0.21 | 5.4±0.90 | μM |
| L-744,832 | Yes | 44±14 | 28±18 | 23±12 | μM |
| FTI-I | Yes | 48±9.7 | 51±5.1 | 52±6.3 | μM |
| FTI-II | Yes | 61±0.79 | 59±8.8 | 58±3.3 | μM |
| GGTI-286 | Yes | 16±3.7 | 50±17 | 56±18 | μM |

Results presented are the mean $IC_{50}$ values ± SEM.

**Supplemental Table 3: The cross-resistance profile of HT1080 EV, HT1080 TMT1A, and HT1080 TMT1B cells with thiol-based GGTIs and DCMB.**

| Drug | HT1080 EV | HT1080 TMT1A | HT1080 TMT1B | Units |
|---|---|---|---|---|
| GGTI-298 | 0.79±0.30 | 4.3±0.17 | 4.7±0.11 | µM |
| GGTI-298 + DCMB | 1.1±0.33 | 1.7±0.34 | 4.3±0.79 | µM |
| GGTI-286 | 21±6.4 | 43±13 | 51±17 | µM |
| GGTI-286 + DCMB | 25±2.5 | 19±2.2 | 66±2.6 | µM |

Results presented are mean $IC_{50}$ values ± SEM.
